# Early cross-coronavirus reactive signatures of protective humoral immunity against COVID-19

**DOI:** 10.1101/2021.05.11.443609

**Authors:** Paulina Kaplonek, Chuangqi Wang, Yannic Bartsch, Stephanie Fischinger, Matthew J. Gorman, Kathryn Bowman, Jaewon Kang, Diana Dayal, Patrick Martin, Radoslaw Nowak, Ching-Lin Hsieh, Jared Feldman, Boris Julg, Eric J. Nilles, Elon R. Musk, Anil S. Menon, Eric S. Fischer, Jason S. McLellan, Aaron Schmidt, Marcia B. Goldberg, Michael Filbin, Nir Hacohen, Douglas A Lauffenburger, Galit Alter

## Abstract

The introduction of vaccines has inspired new hope in the battle against SARS-CoV-2. However, the emergence of viral variants, in the absence of potent antivirals, has left the world struggling with the uncertain nature of this disease. Antibodies currently represent the strongest correlate of immunity against COVID-19, thus we profiled the earliest humoral signatures in a large cohort of severe and asymptomatic COVID-19 individuals. While a SARS-CoV-2-specific immune response evolved rapidly in survivors of COVID-19, non-survivors exhibited blunted and delayed humoral immune evolution, particularly with respect to S2-specific antibody evolution. Given the conservation of S2 across β-coronaviruses, we found the early development of SARS-CoV-2-specific immunity occurred in tandem with pre-existing common β-coronavirus OC43 humoral immunity in survivors, which was selectively also expanded in individuals that develop paucisymptomatic infection. These data point to the importance of cross-coronavirus immunity as a correlate of protection against COVID-19.

## Introduction

The relentless spread and unpredictable nature of disease caused by SARS-CoV-2 continue to paralyze the globe. However, the introduction of potent vaccines has inspired new hope that the end of the pandemic is in sight (1–3). Yet, the slow vaccine rollout, emergence of new viral variants (4–6), confusing results of convalescent plasma trials, incomplete efficacy from monoclonal therapeutics, coupled with the lack of potent antiviral therapeutics, has left the globe with the burden of managing the uncertain nature of this disease. Thus, there is an urgent and continued need to characterize the humoral antibody response to acute disease, and its correlate with outcomes, to better define biomarkers to support clinical care and target the design of monoclonal therapeutics strategies.

SARS-CoV-2 infected patients experience a wide range of clinical manifestations ranging from asymptomatic infection to severe disease that may exacerbate and result in acute respiratory distress syndrome (ARDS) and ultimately death (7). However, while age (8) and comorbidities are enriched in those with more severe disease (9–12), the outcome of SARS-CoV-2 infection is highly variable. Emerging immune correlates analyses have suggested that early robust neutralizing antibody responses (13–15), innate immune responses (16), Fc-effector activity (17, 18), as well as altered B cell and T cell frequencies (19, 20) and viral loads (21–23) have all been linked to differential outcome.

Among these emerging correlates, antibodies have been implicated in both natural resolution of infection (16, 21, 24) as well as protection following vaccination (25, 26) with vaccine antibody-mediated protection observed prior to the evolution of neutralizing antibody activity ((3, 27, 28). Importantly, beyond neutralization, antibodies control or clear infection via their ability to leverage the immune system via their constant domains (Fc), functions that evolve rapidly following natural infection (16). Specifically, antibodies recognizing the S2 domain, the most conserved region of the SARS-CoV-2 spike, evolve most rapidly and predict survival of natural SARS-CoV-2 infection (16). Remarkably, S2-specific neutralizing antibodies have been observed in plasma samples collected prior to the pandemic (29, 30), hypothesized to have emerged in response to common coronavirus infection. However, whether these S2-specific antibodies emerge selectively earliest in infection and track with the presence of common coronavirus immunity remains unclear, and could account for an immunological bias that may temper SARS-CoV-2 infection and further lead to enhanced protective immunity against severe disease.

Thus, using samples from a large acute cohort SARS-CoV-2 infection study (add ref), we profiled the evolving humoral immune response to SARS-CoV-2 over the first twelve days following symptom onset, as well as for a group of asymptomatic individuals. Class switched SARS-CoV-2 S2-specific humoral immune responses evolved rapidly and selectively in acute survivors of COVID-19. Moreover, the presence of a robust common β-coronavirus OC43 response was greatly expanded in individuals that survived infection and experienced milder disease. Most intriguing, these earliest OC43-specific responses were tightly linked to the early evolving SARS-CoV-2 response in survivors of severe and moderate disease, uncovering a unique SARS-CoV-2/OC43 relationship during acute infection that may give rise to the early SARS-CoV-2 response. Moreover, S2-functional humoral immunity was selectively expanded in asymptomatic or paucisymptomatic infections, pointing to the critical nature of these particular humoral immune responses as potential correlates across the disease spectrum. These data point to a critical advantage in eliciting early S2-specific robust functional Fc-effector functions that may emerge from pre-existing β-coronavirus immunity. Thus, rather than original antigenic sin, pre-existing cross-CoV immunity may accelerate the evolution of cross-reactive humoral immune responses to highly conserved regions of the virus, which may point to critical targets of CoV immunity.

## Results

### Dampened SARS-CoV-2 humoral immune evolution is a signature of COVID-19 mortality

Previous studies have noted distinct humoral evolutionary trajectories across individuals with different clinical outcomes following SARS-CoV-2 infection, marked by different magnitudes of humoral immune responses (16, 31), differential targeting of antigens (21), increased functional breadth (31), incomplete IgG class switching (32), or neutralization activity (14). However, the majority of these studies probed humoral immune responses several days to weeks following symptom onset. Thus, to gain insights into the earliest host-pathogen interactions that may underlie differences in the disease trajectory, we profiled a cohort of acutely ill COVID-19 patients. A total of 217 patients with confirmed SARS-CoV-2 infection by nasopharyngeal PCR were collected at the time of admission through the Emergency Department (ED) at approximately 0 - 12 days following symptom onset and stratified by disease severity and 28-day outcome into three groups: (1) moderate - requiring hospitalization and supplemental oxygen support (*n* = 118, corresponding to a max score of 4 on the WHO Ordinal Outcomes scale in 28 days); (2) severe – intubated but survived to 28 days (*n* = 62, corresponding to max WHO scale of 6-7); and (3) deceased - non-survivor (*n* = 37, corresponding to WHO scale 8) (33, 34) **(Fig. 1A, Sup. Table 1)**.

**Figure 1.**
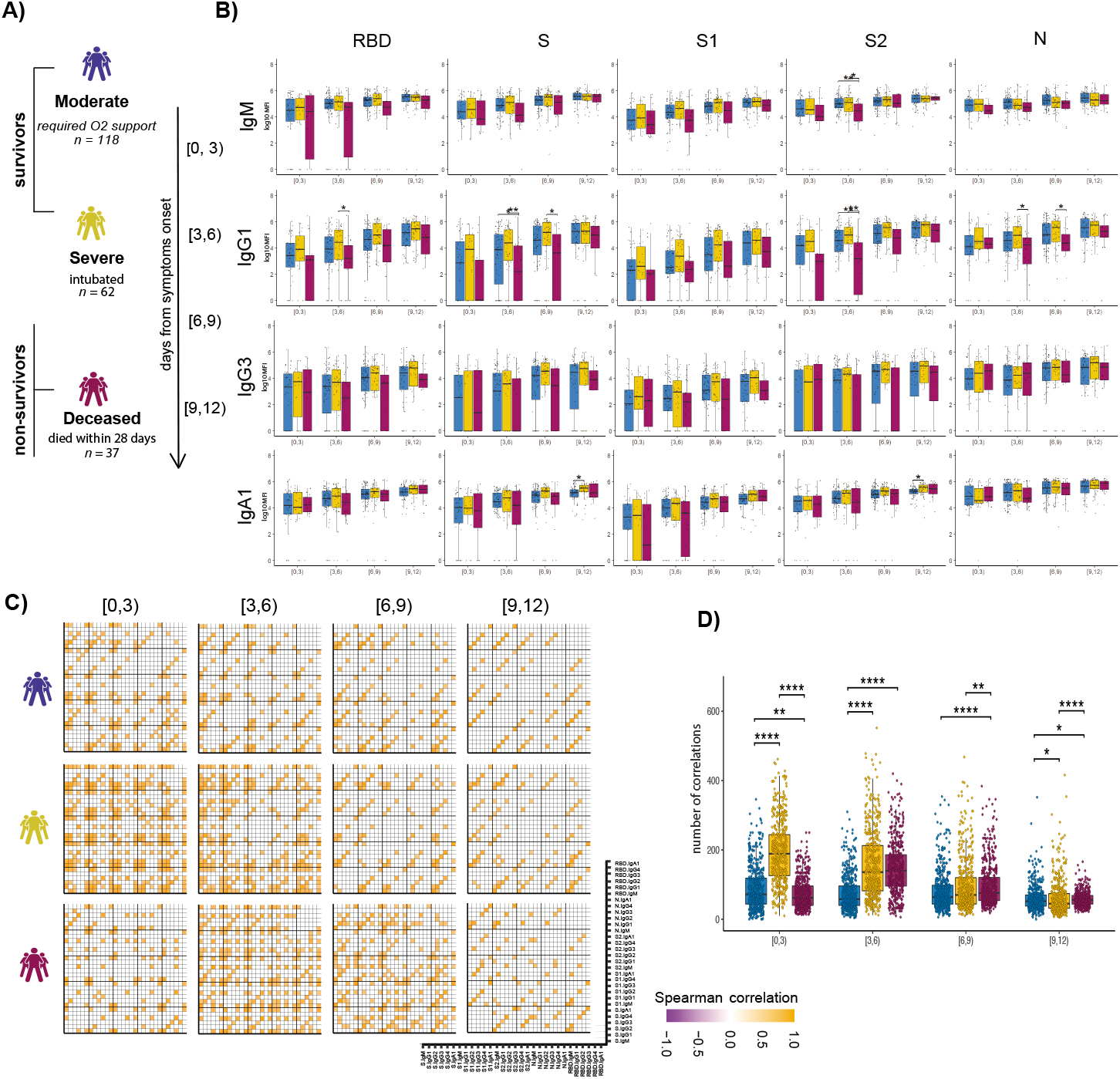
Evolution of early SARS-CoV-2 specific humoral immune responses following symptom onset across acutely ill COVID-19 patients. **(A)** The cartoon shows the study groups based on COVID-19 severity: 217 COVID-19-infected patients were sampled on days 0, 3, and 7 after admission to the hospital. Patients were classified into three groups based on the maximal acuity within 28 days of enrollment: Moderate: hospitalized that required supplemental oxygen (*n* = 118). Severe: intubation, mechanical ventilation, and survival to 28 days (*n* = 62). Deceased: death within 28 days (*n* = 37). Based on the day of symptom onset, the samples were divided into four temporal groups: [0, 3), [3, 6), [6, 9), [9, 12). **(B)** The whisker plots show the distribution of antibody titers across moderate (blue), severe (yellow), and deceased (red) over the study time course. The solid black line represents the median, and the box boundary (upper and below) represents the first and third quartiles. The dots show the scaled values of each sample. A two-sample Wilcox test was used to evaluate statistical differences across groups for all the intervals and features. The P-values were corrected from multiple hypothesis testing using the Benjamini-Hochbery procedure per each interval. Significance corresponds to adjusted P-values. (* p < 0.05, ** p < 0.01). **(C)** The correlation heatmap shows pairwise Spearman correlation matrices of SARS-CoV-2-specific antibody response across COVID-19 severity groups (moderate, severe, and deceased) for all four intervals. Correlation coefficients are shown only if they are larger than 0.6 and statistically significant after Benjamini-Hochberg correction for multiple hypothesis testing. Negative correlations are indicated in purple, positive correlations are shown in orange. (**D**) The statistical evaluation of the effect of sample size. The spearman correlation is calculated by randomly selected ten samples per category for 500 runs. The number of statistically significant correlations (larger than 0.6) is calculated and tested by the Mann-Whitney U test. Significance corresponds to adjusted P-values. (* p < 0.05, ** p < 0.01, *** p <0.001, **** p <0.0001).

System Serological profiling during the acute window of infection pointed to a significant deficit in IgG1 RBD, full S protein, S2, and N-specific antibody, and S2 specific IgM levels in the non-survivor group between 3-9 days after symptom onset **(Fig. 1B)**. Similar trends were noted for SARS-CoV-2-specific IgG3, IgA1 titers, as well as FcγR binding capacity **(Sup. Fig. 1)**. Moreover, while FcγR binding was associated with neutralization in survivors of COVID-19, this relationship was lost in non-survivors (**Sup. Fig. 2**). Given that antibodies are generated as polyclonal swarms, we profiled the coordination of the evolution of the humoral immune response across the groups. Strikingly higher coordination was observed in severe disease survivors within 0-3 days of symptom onset compared to limited coordination in the moderate and non-survivors (**Fig. 1C and 1D**). Additionally, correlations were weak but persisted in the individuals that experienced moderate disease, while they declined over the time for survivors of severe disease. Conversely, coordination in the humoral immune response increased at day 3-6 and immediately decreased in individuals that did not survive SARS-CoV-2 infection. This analysis suggests that severe survivors of COVID-19 generate a more robust, highly coordinated acute humoral immune response very early in disease, compared to patients that succumbed to COVID-19.

### S2-specific responses are selectively enriched in survivors of COVID-19

Considering the multitude of differences across the groups, we next aimed to define whether specific longitudinal humoral features could resolve survivors from non-survivors. Thus, we generated pairs of nested mixed linear models across each set of subjects, accounting for comorbidities, age, and gender, all attributes previously linked to more severe COVID-19 (35) **(Fig. 2A-C)**. First, we compared survivors with severe disease to non-survivors. Following correction for multiple comparisons, six features were selectively enriched among patients that survived COVID-19, four of which were directed at the highly conserved S2-domain of the SARS-CoV-2 Spike **(Fig. 2A)**. Critically, all S2-specific features related to enhanced S2-specific Fc-receptor binding antibodies were highly and selectively enriched among survivors with severe disease compared to non-survivors.

**Figure 2.**
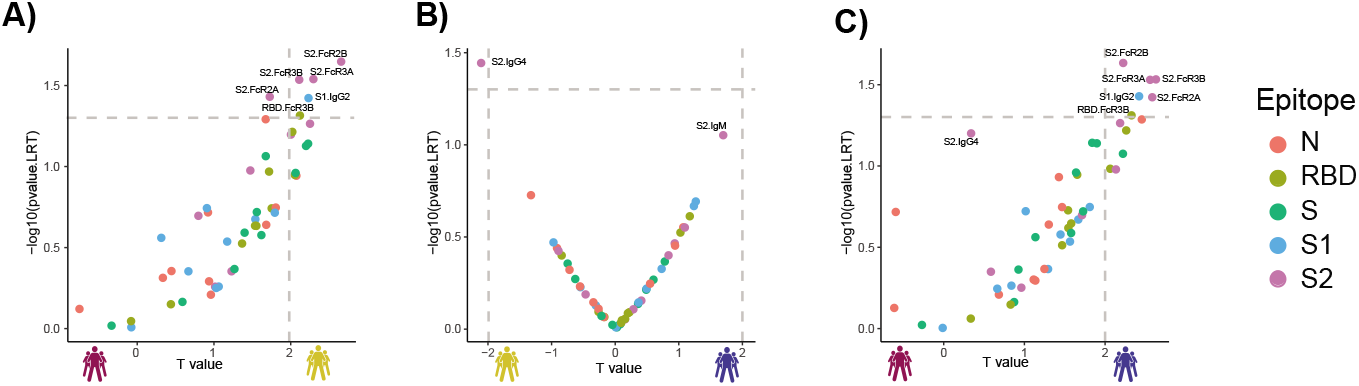
Selective enrichment of S2-specific responses across COVID-19 patients. **(A-C)** Volcano plots of pairwise comparisons across pairs of each of the three groups highlight differences across groups controlling for age, BMI, heart, lung and kidney diseases. The volcano plots include comparisons of **(A)** individuals that passed away within 28 days (deceased) vs. severe survivors; **(B)** subjects who experienced moderate disease vs. severe survivors; **(C)** subjects who ultimately passed away (deceased) vs. subjects who developed moderate disease. The x-axis represents the *t* value of the full model, and the y-axis denotes the *p* values by likelihood ratio test comparing the null model and full model. The null/full model represents the association between each individual measurement (response) and all collected clinical information with/without disease severity (see methods). The horizontal gray dashed line denotes the *p*-value equals 0.05, and the vertical gray dashed line denotes a manually selected threshold (*t* values = 2).

In contrast, a more balanced distribution of distinct antibody features was observed when comparing moderate and severe survivors. Interestingly, a single S2-specific antibody feature, S2-specific IgG4 in severe survivors, was the single statistically significant difference across these groups **(Fig. 2B)**, again illustrating the critical value of S2-specific immunity in resolving disease state. A comparison of survivors with moderate infection compared to non-survivors highlighted the elevated immune responses in individuals with moderate disease, marked by five S2-specific Fc-receptor binding antibody features that reached statistical significance **(Fig. 2C)**. These data highlight the uniquely diverging S2-specific Fc-profiles that represent key early biomarkers that clearly distinguish survivors from non-survivors of COVID-19.

### Common coronavirus responses are enriched in survivors early in infection course

The unconventional patterns of seroconversion in S2-specific immunity in survivors of SARS-CoV-2 infection, marked by a near simultaneous evolution of IgG and IgM at early time points **(Sup. Fig 3)** (13, 22, 36–38), point to the possibility of either remarkably rapid maturation of the humoral immune response, or the potential expansion and evolution of preexisting cross-coronavirus immunity. While common coronaviruses (cCoV) circulate annually, giving rise to broad population level seroprevalence, conflicting data have emerged related to seroprevalence of cCoV-specific immunity among individuals with distinct disease trajectories (39–46). Given the significant prevalence of β-coronavirus OC43 in the USA (47), we profiled the prevalence of OC43 specific-immunity across the groups. To avoid detection of SARS-CoV-2 cross-reactive responses, we focused on the OC43 receptor binding domain (RBD), which is poorly conserved across the β-coronaviruses, offering a unique opportunity to dissect OC43 and SARS-CoV2 immunity. Higher OC43 RBD-specific IgM and IgG1 levels were observed in survivors with severe and moderate disease, particularly within 3-6 days of symptom onset **(Fig. 3A)**. Conversely, no differences were noted in IgA, IgG3, or Fc-receptor binding across the groups. Importantly, only a slight increase was observed in all OC43 responses across the groups over the study period, clearly indicating stability in cCoV immunity that was not boosted by infection. These data point to enrichment of OC43-immunity, but not an evolution of these responses, in survivors in the first week of observation.

**Figure 3.**
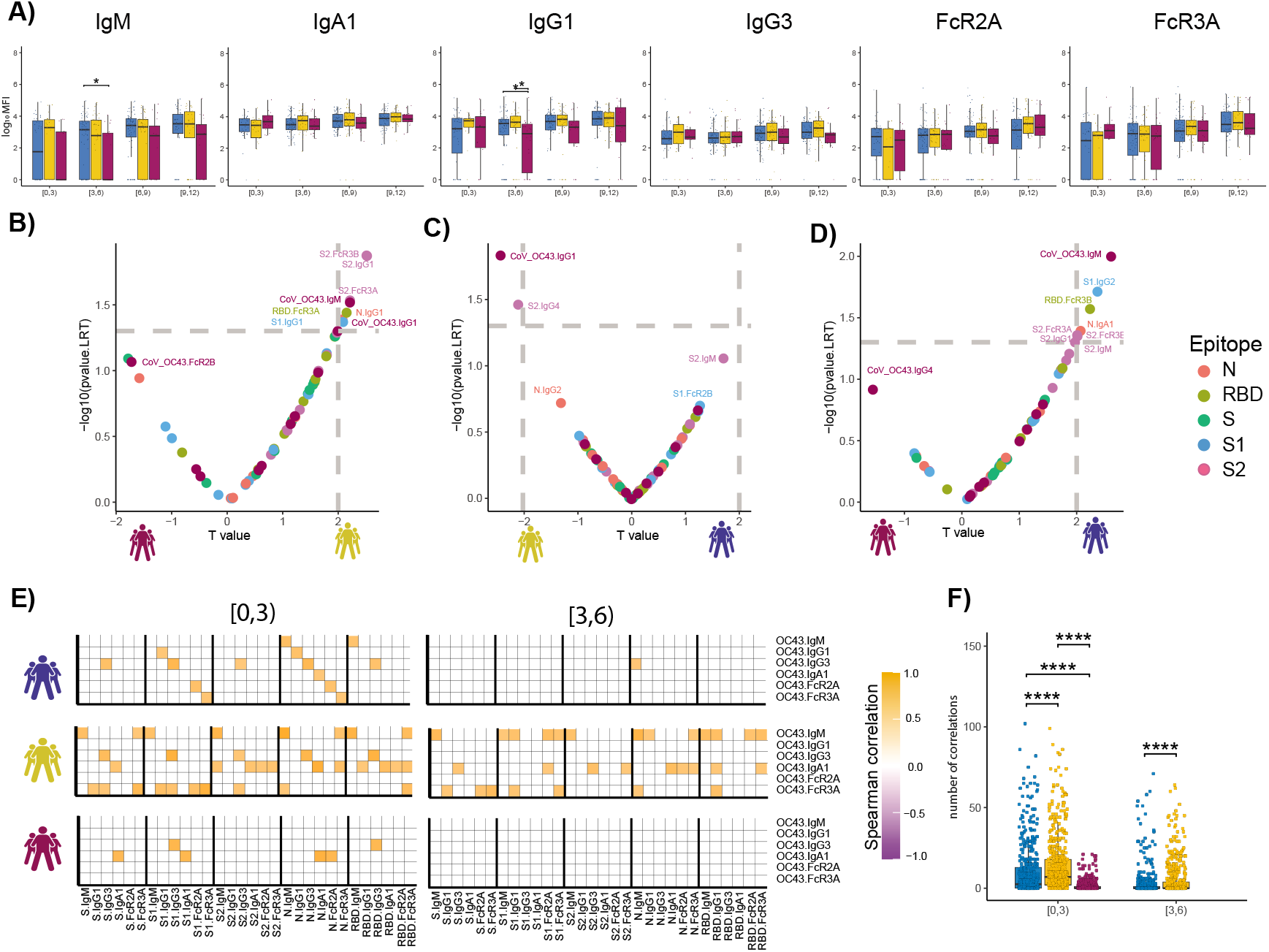
The temporal evolution of the human OC43 specific humoral immune response. **(A)** The whisker bar graphs show the distribution of human OC43 receptor binding domain (RBD)-specific antibody titers and OC43-specific antibody mediated Fc-receptor binding profiles across moderate, severe, and non-survivor COVID-19 groups over the study time course. The solid black line represents the median and box boundary (upper and bottom). **(B-D)** The volcano plots show the pairwise comparisons across the three COVID-19 severity groups, **(A)** individuals that passed away within 28 days (deceased) vs. severe survivors; **(B)** subjects who experienced moderate disease vs. severe survivors; **(C)** subjects who ultimately passed away (deceased) vs. subjects who developed moderate disease, including human OC43 RBD-specific humoral immune data. **(E)** The correlation heatmap shows the pairwise Spearman correlation matrices between OC43-specific and SARS-CoV-2 antibody levels across three COVID-19 severity groups (moderate, severe, and non-survivors) across the study time course. The correlation coefficients were shown only if statistically significant (adjust *p*-value < 0.05) after Benjamini-Hochberg correction from multiple hypothesis testing. **(F)** The statistical evaluation of the effect of sample size. The spearman correlation is calculated by randomly selected ten samples per category for 500 runs (the deceased group in day interval [3,6), is not included since the number of samples is less than 10). The number of statistically significant correlations (larger than 0.6) is calculated and tested by the Mann-Whitney U test. Significance corresponds to adjusted P-values. (* p < 0.05, ** p < 0.01, *** p <0.001, **** p <0.0001).

To begin to define whether the presence of distinct OC43-specific immune responses could resolve the groups, we next integrated OC43 RBD-specific humoral immune profiles into the paired nested mixed linear models. While S2-specific humoral immune responses remained top predictors of severe survivors compared to non-survivors, OC43 RBD-specific IgM antibody levels were significantly and selectively expanded in the survivors (**Fig. 3B**). OC43 RBD-specific IgG1 was selectively expanded in survivors with severe disease compared to moderate disease (**Fig. 3C**). OC43 RBD-IgM was also the most discriminatory feature between individuals with moderate disease and individuals who died within 28 days (**Fig. 3D**). These data point to the unique enrichment of OC43-specific immunity in individuals that selectively and rapidly evolved robust S2-specific immunity linked to the survival of the disease.

To attempt to define whether the presence of cCoV immunity to OC43 could be linked to the evolution of early SARS-CoV-2 immunity differentially across the groups, we next looked at the relationship between the very early OC43- and SARS-CoV-2-response across groups over time, with the assumption that negative correlations would indicate dampened or blocking by pre-existing antibodies and positive correlations may suggest leveraged-cooperativity where the evolution of SARS-CoV-2 responses may emerge from pre-existing OC43 responses. Striking differences were observed in the overall correlational structure of the earliest OC43-specific response with the SARS-CoV-2 response (**Fig. 3E and 3F**). Specifically, positive correlations were observed across OC43/SARS-CoV-2 in survivors compared to non-survivors, with broader correlations in survivors with severe disease, and a more focused OC43-specific IgM, IgG3, IgA, FcR3a with SARS-CoV-2 immunity. Interestingly, relationships were observed across Spike, S1, S2, Nucleocapsid, and the receptor binding domain (RBD), suggesting that the influence of preexisting OC43 resulted in an overall expanded response across SARS-CoV-2 antigens. Conversely, these relationships decayed across all groups just three days later, likely due to an affinity maturation and evolution towards largely SARS-CoV-2 specific immunity over time. However, the fact that these relationships were retained to a greater degree in severe survivors, points to more robust recruitment of these pre-existing responses that may be essential to rapid containment and ultimate clearance of the virus.

### Expanded S2-specific Fc-receptor binding antibodies are selectively expanded in asymptomatic infection

To next test whether these relationships with cCoVs only exist in hospitalized patients, or also extend to protective milder immunity, we next extended this analysis beyond hospitalized patients. Differences in the functional profile of SARS-CoV-2 and OC43-specific antibodies were profiled in a longitudinal community-based cohort of asymptomatic/paucisymptomatic SARS-CoV-2 infected individuals sampled both prior to and after SARS-CoV-2 infection (48). While all individuals harbored robust class switched IgA and IgG OC43 responses, no differences were observed in OC43-specific antibody responses across individuals that evolved no symptoms (level 0), very few symptoms (level 1), or few mild symptoms (level 2) **(Sup. Fig. 4)** prior to or after infection (**Fig. 4A**). Conversely, OC43-specific IgG1 selectively increased following infection, with a more significant expansion among individuals that experienced the fewest symptoms, suggesting that class switched OC43-specific immunity may be leveraged selectively in the containment of asymptomatic/paucisymptomatic SARS-CoV-2 disease.

**Figure 4.**
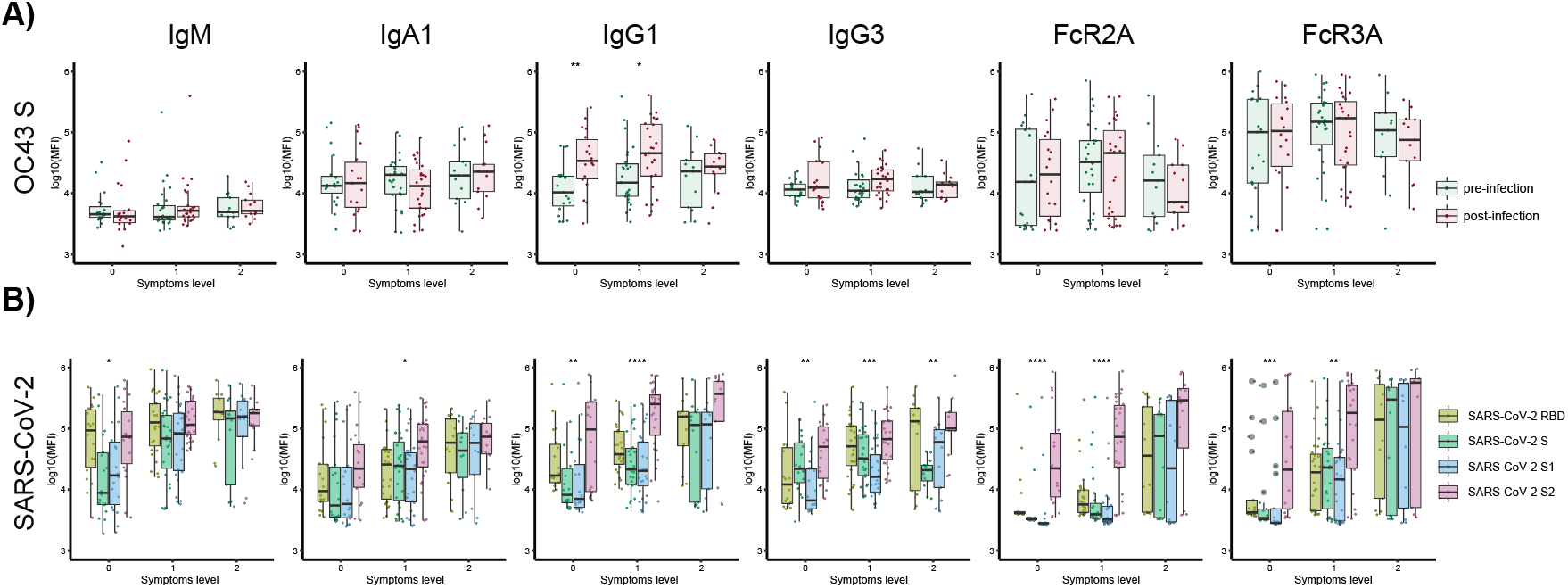
SARS-CoV-2 S2-specific antibody functionality tracks with asymptomatic SARS-CoV-2 infection. **(A)** The whisker box plots show the overall humoral immune response to OC43 RBD-spike titers across a community based SARS-CoV-2 infection cohort divided by individuals that were asymptomatic (symptoms level 0) or experienced symptoms (symptoms level 1 or level 2, based on degree of symptoms) pre- and post-infection. **(B)** The bar graphs illustrate the SARS-CoV-2 specific humoral immune response across the RBD, S, S1, and S2 antigens across the same community based surveillance study divided by the degree of symptoms (symptoms levels). The dots show the scaled values of each sample. A two-sample Wilcox test was used to evaluate statistical differences across different epitopes for all the symptom categories. Significance corresponds to adjusted P-values. (* p < 0.05, ** p < 0.01, *** p <0.001, **** p <0.0001).

We next examined the SARS-CoV-2 specific response. Broad isotype and subclass immune responses (IgM, IgA, IgG1, and IgG3) were noted across the Spike in both asymptomatic and paucisymptomatic (symptoms level 1 and 2) individuals. However, an unexpected difference was noted in the Fc-receptor binding capability of SARS-CoV-2 S2-specific responses. Specifically, individuals with no symptoms or a single symptom elicited solely S2-specific Fc-receptor binding antibodies (**Fig. 4B**) but did not evolve Fc-receptor binding antibodies to other specificities. At a more granular level, while S2-specific IgG1 humoral immune responses were expanded across all individuals with 0-2 symptoms, only S2-specific antibodies were able to interact with the phagocytic FcγR2A and cytotoxic FcγR3A antibodies in asymptomatic individuals. Individuals with one symptom gained more Fc-receptor binding antibodies across antigen specificities, whereas all SARS-CoV-2 specific antibodies had the capacity to interact with Fc-receptors in individuals with at least two symptoms following mild SARS-CoV-2 infection. These data suggest that the expansion of S2-specific functional antibodies may be sufficient to clear asymptomatic SARS-CoV-2 infection, thereby not only representing an immune correlate of protection against death, but also representing a key correlate of protection against symptomatic COVID-19 disease.

## Discussion

In a matter of months, SARS-CoV-2 caused 100s of millions of infections, over a million deaths, and has compromised global economies (49, 50). Public health measures, including masks, distancing, quarantines, have slowed the spread (51, 52). While SARS-CoV-2 specific therapeutics have shown more moderate promise, the use of steroids and ventilation have greatly improved our ability to provide care for individuals that present with severe disease (53). Yet, it is now certain that vaccines will be essential to end this pandemic. However, in the absence of precise correlates of immunity against COVID-19, the development of more potent monoclonal therapeutics and the evaluation of vaccine durability has been difficult.

Humoral immune responses have been linked to both resolution of natural disease (13, 54) and protection following vaccination (1, 55, 56). Specifically, longitudinal studies have reproducibly demonstrated the presence of more robust neutralizing antibody responses among individuals with more severe disease and death (57). By contrast, the ability of antibodies to drive antibody effector functions, via interactions with Fc-receptors, with an early enrichment of S2-specific antibodies among survivors, has been linked to survival (16, 21). However, the precise target of these protective antibodies, as well as the kinetics of their evolution, was unclear. Here, using severely ill acute COVID-19 patients, we profiled the evolution of the earliest antibody response in moderately and severely ill acute COVID-19 patients. Striking early evolution of S2-specific humoral immune response was observed in individuals that survived infection. Specifically, class-switched responses were observed just days after symptom onset, pointing to the emergence of these responses out of memory rather than the de novo induction of these humoral immune profiles. Given the conserved nature of S2 across β-coronaviruses, these data suggest that individuals who survive COVID-19 may have an earlier advantage, able to develop rapid and robust humoral cross-reactive immune responses able to contain and ultimately clear infection.

Unlike the potent neutralizing activity of antibodies against RBD, S2-specific antibodies can neutralize, but are likely to provide protection through additional humoral mechanisms. Interestingly, S2-specific antibodies with Fc-receptor binding capabilities, rather than S2-titers alone, were among the strongest predictors of protective immunity against death. Moreover, S2-specific antibody functions were selectively augmented in asymptomatic SARS-CoV-2 infection, suggesting that the ability of these antibodies to recruit innate immune effector functions may be key to their protective activity. Thus, even in the absence of potent neutralization, it is plausible that functional S2-specific antibodies may play a critical role in the early recognition and clearance of SARS-CoV-2 upon infection. Moreover, given the highly conserved nature of S2, it is plausible that highly functional S2-specific monoclonal antibodies or vaccine-induced immune responses may provide broader and more potent protection against emerging variants of concern and perhaps even additional β-coronaviruses. Furthermore, S2-specific responses exhibit delayed kinetics and magnitude-pointing to an opportunity to develop more potent and functionally enhanced S2-specific monoclonal therapies that may have a broader clinical window of therapeutic opportunity to control/clear the infection.

OC43 is a prevalent β-coronavirus known to cause seasonal colds, like many other CoVs (58). Here we hypothesized that S2-specific humoral immune responses arose from pre-existing responses to OC43 or other common coronaviruses. To test this hypothesis, we examined the early evolution of SARS-CoV-2 specific humoral immunity in relation to the strength of the existing OC43 response. Importantly we did not see any interference- or evidence of original antigenic sin. We observed that not only S2-specific but global SARS-CoV-2 specific humoral immunity was expanded in a coordinated manner with OC43 in survivors-relationships that were subdued in the non-survivors. Because the RBD and S1 share limited homology across the viruses, it is less likely that pre-existing responses to OC43 could directly lead to the evolution of S1- or RBD-specific responses. However, it is plausible that immediately after infection, prior to symptom onset, that the S2-specific response may emerge earliest, followed by a rapid maturation and spreading of the humoral immune response across the viral proteome. The early S2-specific antibodies, in fact, may capture the virus and deliver these antigens to the immune system more aggressively, acting as adjuvants that may drive a highly diverse response across all viral antigens. Thus early, cross-reactive S2-specific antibodies may not only work to provide an early opsonophagocytic clearance/control mechanism but may also act to directly accelerate the induction of immunity to the virus.

Thus collectively, deep humoral profiling of the acute humoral immune response to SARS-CoV-2 pointed to the striking importance of S2-as an early functional humoral target of the protective immune response that may arise from pre-existing common coronavirus immunity. These data clearly indicate a protective, rather than deleterious, role of pre-existing common CoV immunity, that may, in fact, provide a survival advantage and highlights novel therapeutic and vaccine opportunities to drive immunity to regions of the SARS-CoV-2 that are less likely to evolve and that may even provide pan-coronavirus immunity.

## Methods

### Patient cohort and clinical data collection

#### Acutely ill COVID-19 patients

Patients 18 years or older (*n* = 384) with acute respiratory distress and clinical concern for COVID-19 were enrolled in the Emergency Department (ED) in Boston during the peak of the COVID-19 surge (from 3/24/2020 to 4/30/2020), 306 of whom tested positive for SARS-CoV-2 nasopharyngeal PCR as described by Filbin et al. (34). This analysis includes SARS-CoV-2-positive patients whose symptoms onset was between 0-12 days prior to presentation and whose illness severity and 28-day outcome were classified into three groups; (1) moderate – hospitalized and requiring oxygen support but not mechanical ventilation (*n* = 118, corresponding to WHO Ordinal Outcomes scale 4); (2) severe – intubated but survived to 28 days (*n* = 62, corresponding to WHO scale 6-7); and (3) deceased within 28 days - nonsurvivor (*n* = 37, corresponding to WHO scale 8) (33). Of the 42 COVID-19 patients who died, 24 (57%) received mechanical ventilation, and 18 (43%) did not. Patients were excluded from analysis if they were discharged directly from the ED and were not hospitalized within the next 28 days, or if they were admitted but did not require supplemental oxygen, given these either had mild disease or lacked pulmonary manifestations of COVID-19.

Day 0 blood samples were obtained with the initial clinical blood draw in the ED, and day three and day seven samples were obtained during patients’ hospitalization. The clinical course was followed 28 days post-enrollment to establish outcomes. Samples and clinical information were collected according to an institutional IRB-approved protocol (34). Symptom duration upon presentation was obtained via chart review. Demographic, medical history, and clinical data were collected and summarized for each outcome group, using medians with interquartile ranges and proportions with 95% confidence intervals, where appropriate.

#### Community-acquired mild and asymptomatic COVID-19 individuals

Industry employees (Space Exploration Technologies Corp.) were volunteer tested for COVID-19 starting in mid-April 2020. All employees were invited to participate by email, and there were no exclusion criteria. Participants completed a study survey including the collection of COVID-19 related symptoms (48). Upon obtaining informed consent, blood samples were collected approximately every 39.7 days (standard deviation 13.8 days). Symptoms were classified by severity, with 2 points being assigned to loss of smell/taste, fever, feverish/chills, or cough, and 1 point being assigned to other symptoms such as increased fatigue, headache, congestion, nausea/vomiting, diarrhea, sore throat, and body/muscle aches. Symptom scores were summed and each subject was categorized into one of three levels based on degree of symptoms: level 0 (*n* = 18), no symptoms; level 1 (*n* = 27), mild, symptom score 1-5; level 2 (*n* = 13), moderate, symptom score 6-14. The study protocol was approved by the Western Institutional Review Board. The use of de-identified data and biological samples was approved by the Mass General Brigham Healthcare (previously Partners Healthcare) Institutional Review Board. All participants provided written informed consent.

### Luminex

SARS-CoV-2 and eCoV-specific antibody subclass/isotype and Fcγ-receptor (FcγR) binding levels were assessed using a 384-well based customized multiplexed Luminex assay, as previously described (59). SARS-CoV-2 receptor binding domain (RBD) (kindly provided by Aaron Schmidt, Ragon Institute), SARS-CoV-2 nucleocapsid (N) protein (Aalto BioReagents), and SARS-CoV-2 spike protein (S) (kindly provided by Eric Fischer, Dana Farber), SARS-CoV-2 subunit 1 and 2 of the spike protein (S1 and S2) (Sino Biological), as well as human eCoV antigens: hCoV-OC43 RBD (kindly provided by Aaron Schmidt, Ragon Institute), hCoV-OC43 spike protein (S) (Sino Biological), hCoV-HKU1 spike protein (S) (Immune Tech), SARS-CoV-1, MERS spike proteins (S) (kindly provided by Jason McLellan, University of Texas) were used to profile specific humoral immune response. A mix of HA A/Michigan/45/2015 (H1N1), HA A/Singapore/INFIMH-16-0019/2016 (H3N2), B/Phuket/3073/2013 (Immunetech) was used as a control. Antigens were coupled to magnetic Luminex beads (Luminex Corp) by carbodiimide-NHS ester-coupling (Thermo Fisher). Antigen-coupled microspheres were washed and incubated with plasma samples at an appropriate sample dilution (1:500 for IgG1 and all Fcγ-receptors, and 1:100 for all other readouts) for 2 hours at 37°C in 384-well plates (Greiner Bio-One). Unbound antibodies were washed away, and antigen-bound antibodies were detected by using a PE-coupled detection antibody for each subclass and isotype (IgG1, IgG2, IgG3, IgG4, IgA1, and IgM; Southern Biotech), and Fcγ-receptors were fluorescently labeled with PE before addition to immune complexes (FcγR2A, FcγR2B, FcγR3A, FcγR3B; Duke Protein Production facility). After 1h incubation, plates were washed, and flow cytometry was performed with an IQue (Intellicyt), and analysis was performed on IntelliCyt ForeCyt (v8.1). PE median fluorescent intensity (MFI) is reported as a readout for antigen-specific antibody titers.

### Quantification and Statistical Analysis

All analyses were performed using R version 4.0.3. All the figures were created with many R-supported packages, mainly including ggplot, ggrepel, ggpubr.

### Data Pre-processing

The raw MFI was scaled by the log10 function and then was subtracted by the corresponding PBS values. The normalized MFI values were assigned to zero if they were negative.

### Univariate Plots

The box-plot summarizes the median value with first and third quantiles of clinical groups (moderate, severe, and deceased) across day ranges from symptom onset with the interval of three days (day ranges 0 – 3, 3 – 6, 6 – 9, 9 – 12). The paired *p*-value was estimated by the Mann-Whitney U test for each sub-feature on the individual temporal course across different clinical groups and adjusted by the Benjamini-Hochbery procedure of multiple testing correction. The visualization was performed by the function ‘ggplot’ of R package ‘ggplot2’ (3.3.3), and the p-value was estimated by the function ‘wilcox_test’ and ‘adjust_pvalue’ in the R package ‘rstatix’(0.6.0) and labeled by the function ‘stat_pvalue_manual’ in the R packge ‘ggpubr’ (0.4.0).

### Correlation Analysis

Spearman correlation was used to evaluate the relationship between different measurements and performed by the function ‘rcorr’ of R package ‘Hmisc’ (4.4.2). The time-specific correlation analysis between different antibody measurements was used to exploit the temporal coordination. The Spearman coefficient between titer values at t1 and t2 across the samples in clinical groups respectively were calculated, where t1 and t2 were the defined time courses with the interval of three days. The significance of correlation was adjusted by the Benjamini-Hochbery procedure of multiple testing correction.

In order to evaluate the effect of different sample sizes across disease severities and time intervals, two strategies were applied here. Firstly, for the purpose of visualization, correlation coefficient was considered only if it was larger than 0.6 since the small coefficient may be significant if the number of samples is large. Furthermore, the downsampling strategy was applied to make the size of samples the same across different groups for statistical evaluation. In detail, we randomly downsampled ten samples and calculated the Spearman correlation for 500 runs. Then, the number of significant correlations larger than 0.6 was calculated and tested by the Mann-Whitney U test and multiple testing correction.

### Association analysis between measured antibody levels and clinical outcome by controlling potential cofounders

We accessed the significance of the association between measured antibody levels and clinical outcome by controlling collected potential cofounders implemented by two nested mixed linear models (null and full model) without/with clinical outcomes. We fit two linear mixed models and estimated the improvement in model fit by likelihood ratio test to assess how many measurements have a significantly better fit with the full model at threshold < 0.05.

Null model:

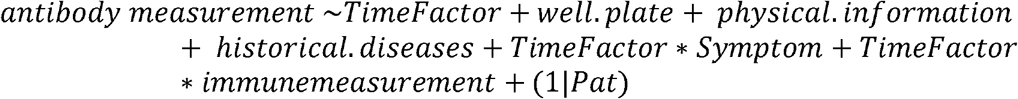

Full model:

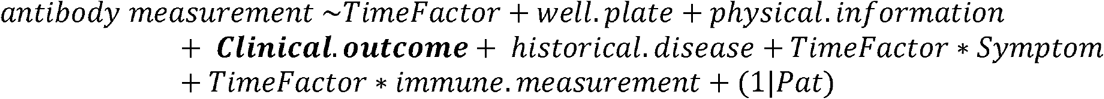

Likelihood Ratio Test: 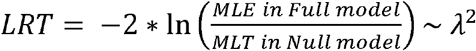

Here, the *historical.diseases* include comorbidities such as heart, lung, kidney, and diabetes, while *physical.information* include age and BMI categories. *Well.plate* represents the batch indicator of Luminex platform. *Time-related* symptoms category include respiratory symptom, fever, gastrointestinal symptoms, and other immune measurements, such as absolute neutrophil count, absolute monocyte count, creatinine, the level of C-reactive protein (CRP), d-dimer, lactate dehydrogenase (LDH) along the time, and troponin level at 72 hours. The R package ‘lme4’ was used to fit the linear mixed model to each measurement and test for measurement across the contrast of interest. The *p*-value from the Likelihood Ratio test and t value of Clinical outcome in Full model was visualized as volcano plot using the ‘ggplot’ function in R package ‘ggplot2’.

## Code availability

All code is publicly available, and the source is indicated in the text and/or the Methods section. Scripts will be made available upon reasonable request. Source data are provided with this paper.

## Acknowledgments

We thank Nancy Zimmerman, Mark and Lisa Schwartz, an anonymous donor (financial support), Terry and Susan Ragon, and the SAMANA Kay MGH Research Scholars award for their support. We acknowledge support from the Ragon Institute of MGH, MIT and Harvard, the Massachusetts Consortium on Pathogen Readiness (MassCPR), the NIH (3R37AI080289-11S1, R01AI146785, U19AI42790-01, U19AI135995-02, U19AI42790-01, 1U01CA260476 – 01, CIVIC75N93019C00052), the Gates Foundation Global Health Vaccine Accelerator Platform funding (OPP1146996 and INV-001650), and the Musk Foundation. MBG, MRF, and NH received support from the American Lung Association and the MGH Executive Committee on Research. This work was also supported by the Translational Research Institute for Space Health through NASA Cooperative Agreement NNX16AO69A

## Contributions

P.K., C.W., D.A.L., and G.A. analyzed and interpreted the data. P.K., S.F., Y.C.B., M.J.G., K.B., and J.K. performed experiments. C.W. and D.A.L performed the analysis. M.G., M.F., N.H., and D. D. managed samples and data collection for acutely ill COVID-19 patients. P.M., A.S.M., E.J.N. and E.R.M. managed samples and data collection for the Community-acquired COVID-19 cohort. R.M., C.H., J.F., A.S., and J.S.M., E.S.F. produced SARS-CoV-2 and OC43 antigens. G.A. and D.A.L. supervised the project. P.K., G.A., and C.W. drafted the manuscript. All authors critically reviewed the manuscript.

## Supplementary Figures

**Supplement Figure 1. Distribution of RBD-, S-, S1-, S2- and N-specific antibody isotypes/ subclasses and their ability to bind Fc-receptors across acutely ill COVID-19 patients.**

The immune response of SARS-CoV-2 infected individuals was profiled against SARS-CoV-2 antigens. Distributions of IgG2, IgG4 as antibody titers and binding to FcγR2A, FcγR2B, FcγR3A, FcγR3B across moderate (blue), severe (yellow) and deceased (red) COVID-19 patients are shown as box-plots over the time intervals (0-3, 3-6, 6-9, 9-12) following symptom onset. The solid black line represents the median, and the upper and bottom lines of box plots show the first and third quartiles. Values are reported as log10 MFI. A two-sample Wilcox test was used to evaluate statistical differences across groups for all the intervals and features. The P-values were corrected from multiple hypothesis testing using the Benjamini-Hochberg procedure per each interval. Here, significance corresponds to the adjusted P-values. (* *p* < 0.05, ** *p* < 0.01).

**Supplement Figure 2. The Spearman correlation between SARS-CoV-2 and human cCoV OC43 antibody levels and neutralization level over time**. **(A)** The correlation heatmap between isotypes/subclasses and neutralization level across different SARS-CoV-2 epitopes measured by Spearman Correlation. **(B)** The correlation heatmap between OC43 -specific isotypes/subclasses and neutralization across different time courses. The adjusted *p* values by the Benjamini-Hochberg procedure were labeled as asterisk (\**p* < 0.05, ** *p* < 0.01, *** *p* < 0.001).

**Supplement Figure 3. Temporal evolution of SARS-CoV-2 specific antibody. (A)** Distributions of IgM, IgG1, IgG3, and IgA1 ratios among multiple SARS-CoV-2 specific antibodies across the different time course. A two-sample Wilcox test corrected from multiple hypothesis testing using the Benjamini-Hochberg procedure pre-time course was used to evaluate statistical differences. Significance represents the adjusted P-values: (* p < 0.05, ** p < 0.01, *** p < 0.001).

**Supplement Figure 4. Defined symptom groups for community acquired-COVID-19 mild and asymptomatic individuals.** The patients were divided into groups with or without COVID-19 symptoms based on 14 symptoms reported in a questionnaire. More severe COVID-19 symptoms, such as loss of smell/taste, fever, feverish/chills, or cough, were scored higher (2 points), and other, such as increased fatigue, headache, congestion, nausea/vomiting, diarrhea, sore throat, and body/muscle aches, were score with 1 point. The scale shows weighted numbers of observed symptoms and is visualized from the top (asymptomatic group: red) to bottom (an asymptomatic group with an increased number of symptoms: blue to red). **(A)** The symptom heatmap. Patients with reported clinical symptoms were marked in red, lack of symptoms was reported in blue. **(B)** The counts of relationships between defined symptoms and groups defined by the titer difference between pre-existing OC43 S-IgG1 and its corresponding titer value after SARS-CoV-2 infection.

**Supplement Table 1. Samples distribution across disease severity and days intervals since symptoms onset.**

