## Supplemental Table and Figures for "Early cross-coronavirus reactive signatures of protective humoral immunity against COVID-19"

| #samples/#patients | deceased | severe | moderate | Total |
|---|---|---|---|---|
| **day 0-3** | 14/14 | 21/21 | 43/43 | 78/78 |
| **day 3-6** | 22/22 | 42/42 | 77/77 | 141/141 |
| **day 6-9** | 17/17 | 50/50 | 76/76 | 143/143 |
| **day 9-12** | 17/17 | 38/38 | 52/52 | 107/107 |
| **Total** | 70/37 | 151/62 | 248/118 | 469/217 |

Supplemental Table

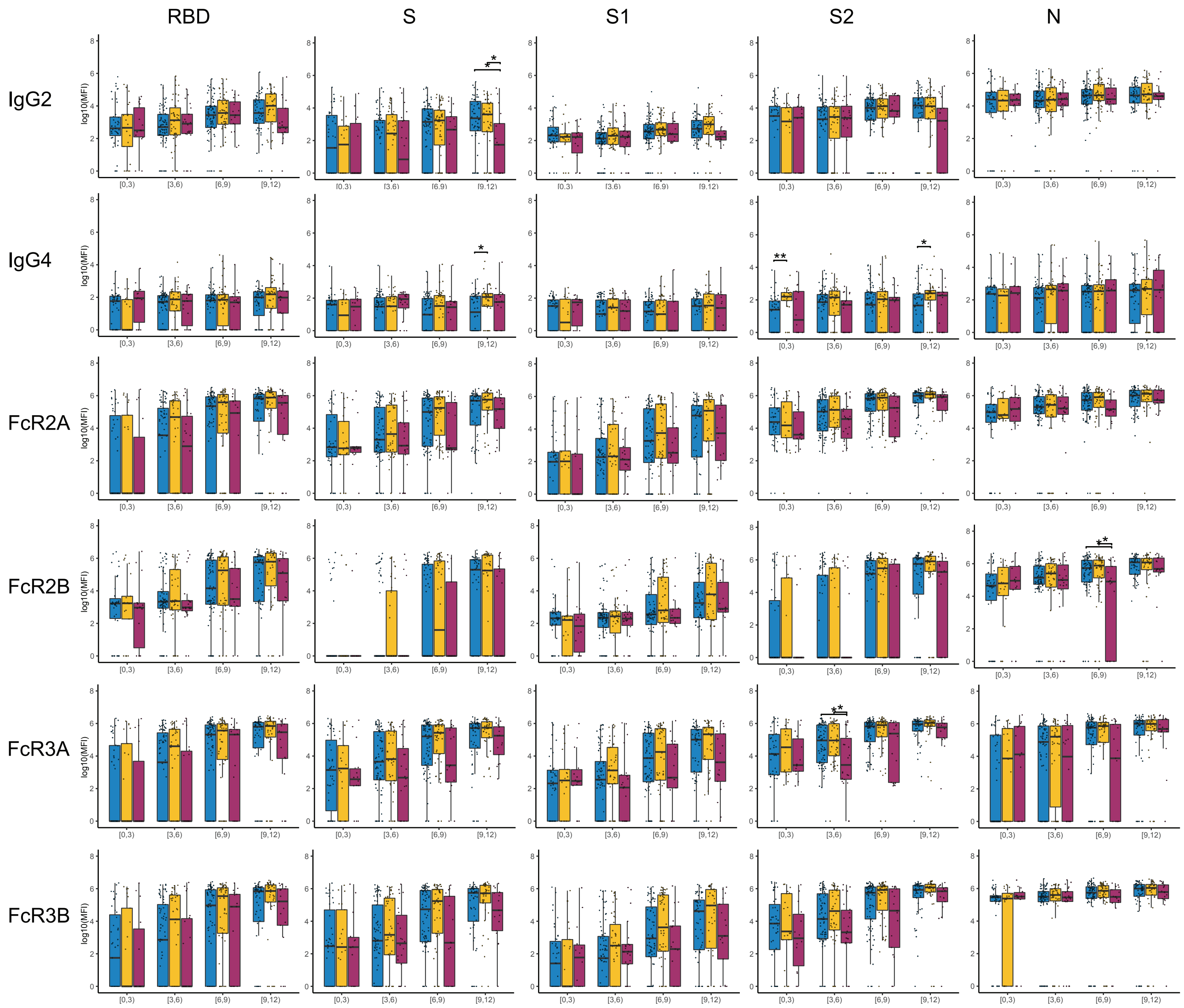


Supplemental Figure 1

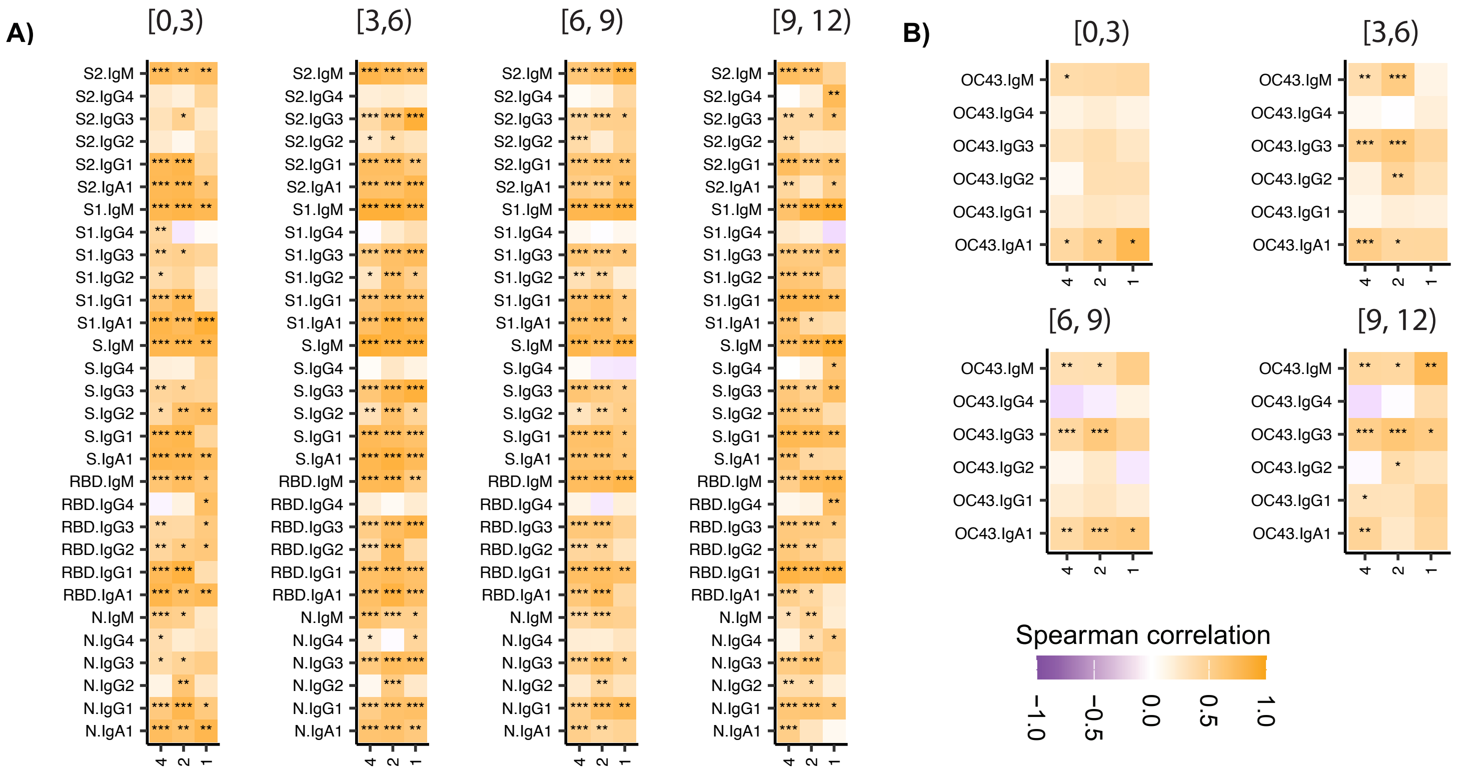


Supplemental Figure 2

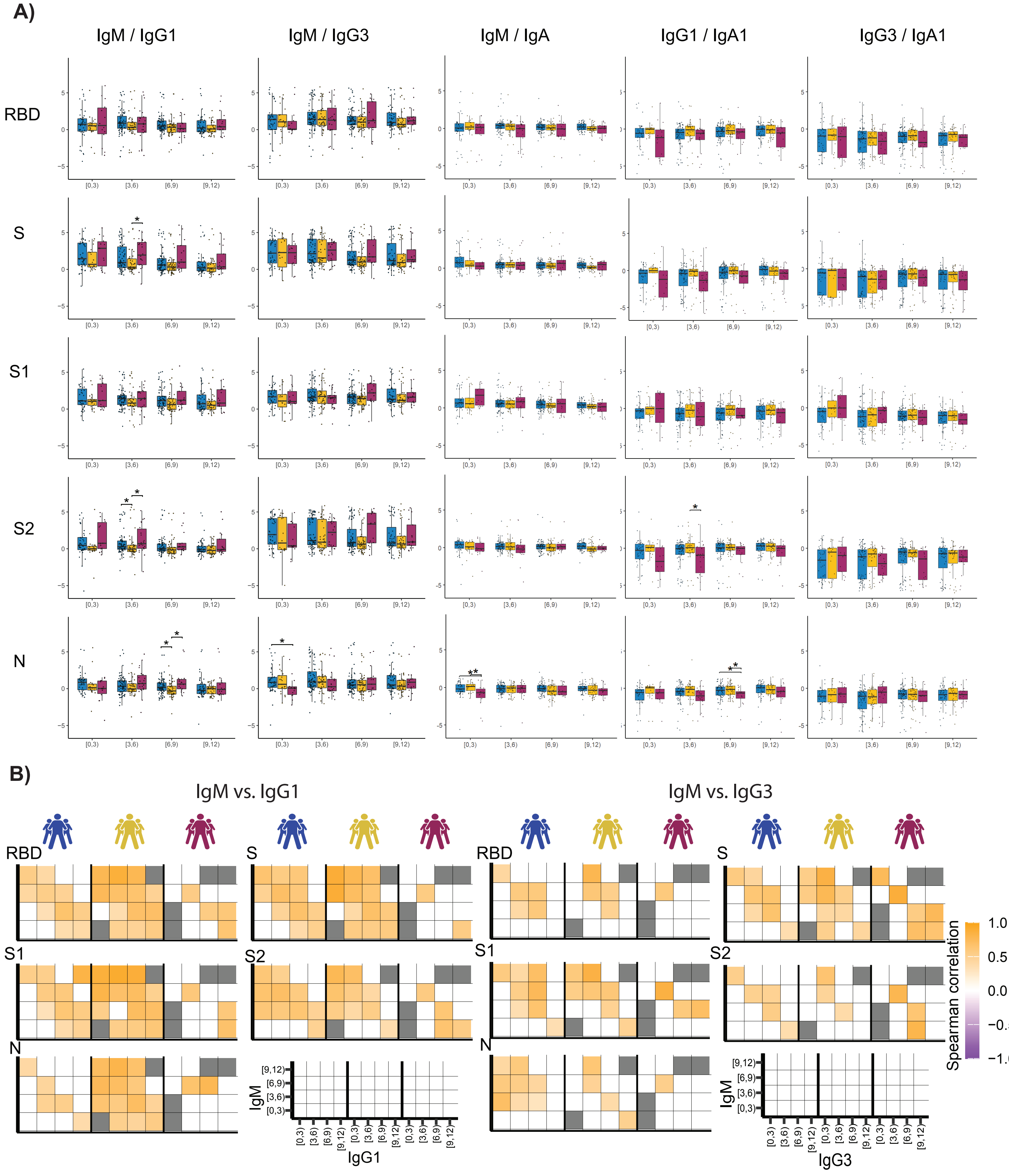


Supplemental Figure 3

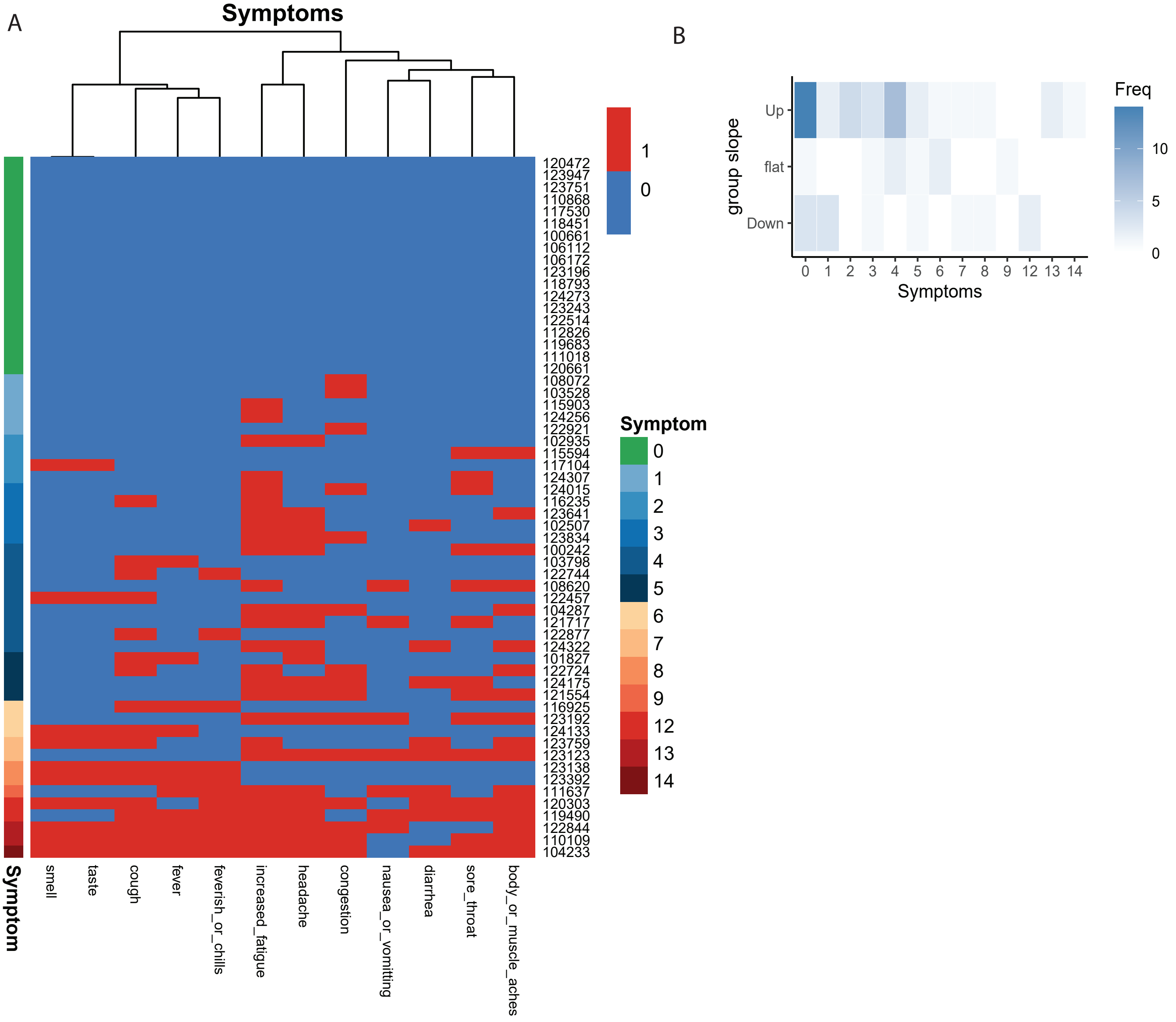


Supplemental Figure 4
